# Engineering Protein Activity into Off-the-Shelf DNA Devices

**DOI:** 10.1101/2022.01.03.474821

**Authors:** Harsimranjit Sekhon, Stewart N. Loh

**Affiliations:** Department of Biochemistry and Molecular Biology, SUNY Upstate Medical University, Syracuse, NY 13210

**Keywords:** Protein Engineering, Alternate frame folding, Toehold mediated strand displacement, DNA logic gates, DNA aptamers

## Abstract

DNA-based devices are relatively straightforward to design by virtue of their predictable folding, but they lack biological activity. Conversely, protein-based devices offer a myriad of biological functions but are much more difficult to design due to their complex folding. This study bridges the fields of DNA engineering and protein engineering to generate a protein switch that is activated by a specific DNA sequence. A single protein switch, engineered from nanoluciferase using the alternate frame folding mechanism and herein called nLuc-AFF, is paired with different DNA technologies to create a biosensor for a DNA or RNA sequence of choice, sensors for serotonin and ATP, and a computational device that processes two DNA inputs. nLuc-AFF is a genetically-encoded, ratiometric, blue/green-luminescent biosensor whose output can be quantified by cell phone camera. nLuc-AFF is not falsely activated by decoy DNA and it retains full ratiometric readout in 100 % serum. The design approach can be applied to other proteins and enzymes to convert them into DNA-activated switches.

## Introduction

Structural biology of proteins and nucleic acids has given rise to a diverse set of methods for biomolecular engineering. DNA building blocks have been used to create self-assembling nanostructures^1,2,3,4^, tools for the detection of other biomolecules^5,6,7^, and ribocomputing devices that control gene expression based on various inputs^8,9^. *De novo* protein design and modifications of existing proteins have also spawned a toolbox of biotechnologies for use in molecular detection^10,11,^, allosteric^12,13^ and optogenetic^14^ control of biological activity, and many more applications. Nucleic acid switches offer the benefit of being inherently easy to design due to their largely predictable folding, but they have limited applications due to their lack of biological activity. Proteins offer the advantage of being biologically active, but they are much more difficult to design *de novo*, and modifying natural proteins often produces unpredictable results. The fields of nucleic acid and protein engineering have largely evolved independently, with little overlap between the two. In this study, we bridge these disciplines to give rise to a novel class of protein switches that can be activated by a DNA input. We use well-developed DNA engineering techniques to activate a color-change ratiometric luminescent biosensor constructed using the alternative frame folding (AFF) mechanism.

AFF is a protein engineering approach that introduces allostery into a protein that has none^15^. The target protein is constructed such that it can adopt either its native fold or one corresponding to a circular permutant (CP). The ability to fold into one of two ‘frames’ in a mutually exclusive fashion is enabled by duplicating an N-terminal (or C-terminal) segment of the protein and fusing it to the opposite end of the WT protein, using a linker peptide long enough to span the distance between the protein’s original N- and C-termini (Fig. 1A). The central, shared segment can then fold with either of the duplicated segments but not both. The populations of protein in native and the CP folds are determined by the relative stabilities of each fold. This property can be used to construct a biosensor that has a different activity in each of the two folds, either through mutation or fusion with other proteins. The relative stabilities of the two folds can then be adjusted in response to a ligand by inserting a disordered ligand binding domain into one of the frames. Provided that the binding domain folds upon encountering the ligand and has a reasonably long N-to-C terminal distance (>20 Å) when folded, this reaction stretches the structure of the fold in which its resident, causing the target protein to switch to the other folding frame.

**Figure 1.**
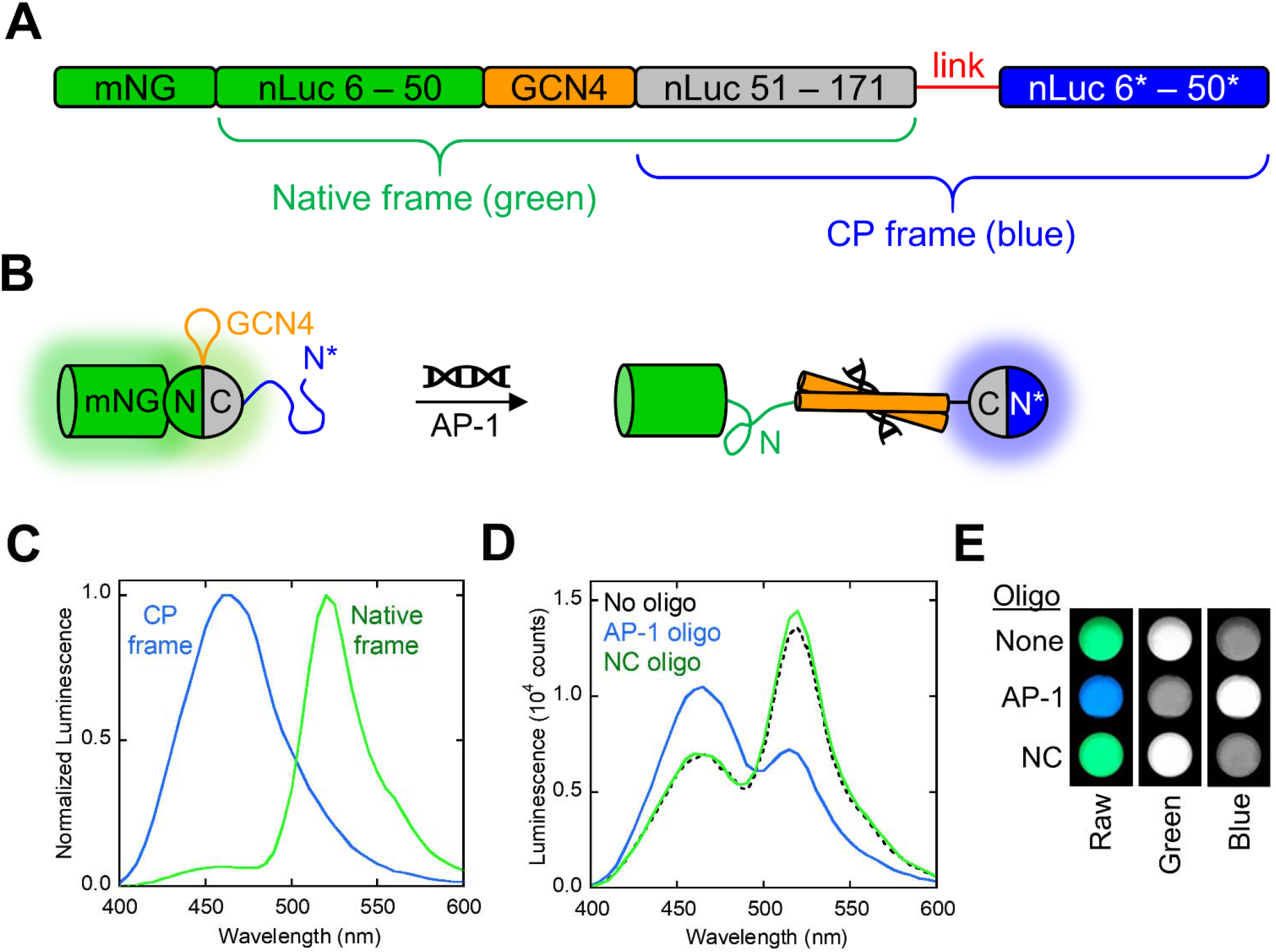
Design and characterization of the nLuc-AFF switch. **(A)** The GCN4 binding domain (orange) was inserted between residues 50 – 51 of the nLuc domain of the GeNL protein (green and grey) to create the native frame of the switch (green-orange-grey). To establish the CP frame of the switch (grey-red-blue), the segment of nLuc that was N-terminal to GCN4 (residues 6 – 50) was then duplicated (residues 6* – 50*; blue) and appended to the C-terminus using a 6-AA linker (red). nLuc-AFF can fold either in the native frame (green luminescence) or the CP frame (blue luminescence) but not both. **(B)** The green-to-blue luminescence change is triggered by DNA binding-induced folding of the GCN4 domain, which stretches and unfolds the native frame and forces nLuc-AFF to fold in the CP frame. This fold shift separates the mNG and nLuc domains and causes luminescence to change from green to blue. Only one monomer of the GCN4 dimer is shown for clarity. **(C)** The isolated native and CP frames exhibited the expected green and blue luminescence spectra, respectively. **(D)** nLuc-AFF emitted mostly green light in the absence of DNA as well as in the presence of NC DNA, and emitted mostly blue light in the presence of AP-1 DNA. **(E)** The green-to-blue color change upon AP-1 binding was visible in raw cell phone pictures as well as by isolating individual green and blue channels from phone images.

Our goal was to make a luminescent biosensor that changes from green to blue emission in the presence of a target DNA sequence. We chose to base the AFF scaffold on nanoluciferase (nLuc, a blue luminescent protein derived from deep-sea shrimp *Oplophorus gracilirostris*) because it possesses high thermodynamic stability^16^ (which helps it retain function after ligand binding domain insertion and circular permutation) and is the brightest luciferase currently known^17^. To establish the blue and green states of the sensor, we employed the variant of nLuc (GeNL) in which the GFP variant mNeonGreen (mNG) was fused to the N-terminus of nLuc^18^. The blue emission of nLuc is captured by mNG and essentially completely converted to green fluorescence via bioluminescent resonance energy transfer. For efficient resonant coupling, it was essential to bring the two proteins into proximity by deleting the first four amino acids of nLuc and the last ten amino acids of mNG^18^.

Our hypothesis was that a green/blue luminescent switch could be engineered by introducing an intramolecular conformational change that extinguishes luminescence of the nLuc domain fused to mNG and turns on luminescence of a second, circularly-permuted nLuc within the same molecule, via the AFF mechanism. As a result of the AFF conformational change, permuted nLuc becomes separated from mNG by a >100 amino acid segment, the length of which is predicted to be >75 Å (*vide infra*). Permuted nLuc is thus expected to emit blue luminescence (Fig. 1B). To couple this conformational change to DNA binding, we turned to GCN4 DNA binding domain as the input domain. GCN4 is a 56 amino-acid peptide consisting of a constitutively dimerized, C-terminal leucine zipper and an N-terminal DNA binding region that is largely unstructured in the absence of its 11-nucleotide consensus DNA sequence (AP-1). GCN4 folds into a rigid, rod-shaped molecule of 75 Å length when bound to AP-1^19^ (Fig. 1B). We previously applied this folding reaction to stretch and unfold the enzyme barnase in the GCN4-barnase fusion protein^20^. We incorporate the AP-1 sequence into off-the-shelf DNA devices to construct two-input logic gates as well as biosensors for ATP and serotonin.

## Results

### Development of the nLuc-AFF biosensor

Beginning with the GeNL protein, we duplicated the N-terminus of nLuc (residues 1 – 50; the duplicated sequence is denoted by asterisks) and appended residues 1* – 50* to the C-terminus of the molecule using a 12-AA linker to generate the nLuc-AFF scaffold. The native and CP frames are shown in Fig. 1A. We chose to duplicate this segment because position 50 is at a surface loop which was previously shown to tolerate circular permutation^21^ as well as insertions^18^. To couple the native-to-CP fold shift to DNA binding, we inserted GCN4 in the surface loop at position 50 of the native fold. In the ligand-free (OFF) state of the switch, GCN4 is predicted to resemble an extended, disordered surface loop in the absence of AP-1. The native fold should tolerate this insertion well and the OFF state is expected to be green. In the AP-1-bound (ON) state, the length of GCN4 will stretch the 6.3 Å C_α_-C_α_ distance between the ends of the loop, splitting the native frame and triggering the fold shift to the CP frame (Fig. 1B). The CP form of nLuc is separated from mNG by 118 amino acids, 56 of which are a 75 Å rod, and the nLuc-AFF is thus expected to emit blue light in the CP frame. We constructed and purified the individual native and CP folds and verified that they were functional green and blue luminescent proteins, respectively (Fig. 1C).

The first iteration of nLuc-AFF exhibited mostly blue luminescence, indicating that it was functional but already in the ON state in the absence of AP-1 (Supporting Fig. S1). We consequently destabilized the CP frame by shortening the linker that connected the N- and C-termini of the CP frame from 12 AA to 8, 6, 4, and 2 AA. Shorter linkers can destabilize a permuted protein by forcing its termini together, a phenomenon previously demonstrated with barnase^22^. Shortening the linkers progressively lowered the blue:green ratio as expected, but the population of CP frame was still too large even at 2 AA linker length (Supporting Fig. S2A).

To further destabilize the CP fold, we deleted residues 1* – 4* from the CP frame (VFTL; the same residues that had been removed from the native frame to make the GeNL variant). We then joined the termini of the CP frame with 10, 6, and 4 AA (with the VFTL truncation the effective linker lengths were 6, 2, and 0 AA). The blue:green ratio diminished with decreasing linker length as before (Supporting Fig. S2B), and this ratio increased after addition of AP-1 for all three variants (Supporting Fig. S3B). This increase was largely due to a decrease in green luminescence without an increase in blue luminescence for the 0 AA linker construct. The 6 AA linker variant had the best combination of high ratiometric change on AP-1 binding and low background signal, and we used this construct for all further experiments. These results demonstrate that the blue/green populations of the nLuc-AFF sensor can be rationally tuned by adjusting linker length in the CP frame.

### nLuc-AFF retains the affinity of input GCN4 domain with its consensus oligonucleotide

The apparent affinity for the nLuc-AFF for AP-1 was measured by adding various concentrations of AP-1 to a fixed amount of biosensor and measuring the ratiometric change in blue:green emission using a scanning plate reader (dividing intensity at 460 nm by intensity at 520 nm; denoted as L460/L520) or cell phone camera (dividing the intensity of the blue channel by the intensity of the green channel). The apparent K_D_ of 12.3 nM ± 5.2 nM (Fig. 2A, Supporting Fig. S4A) is similar to the reported affinity of the isolated GCN4 peptide for AP-1 (22 nM ± 9 nM)^23^, indicating that the GCN4 domain retained full DNA binding activity in nLuc-AFF.

**Figure 2.**
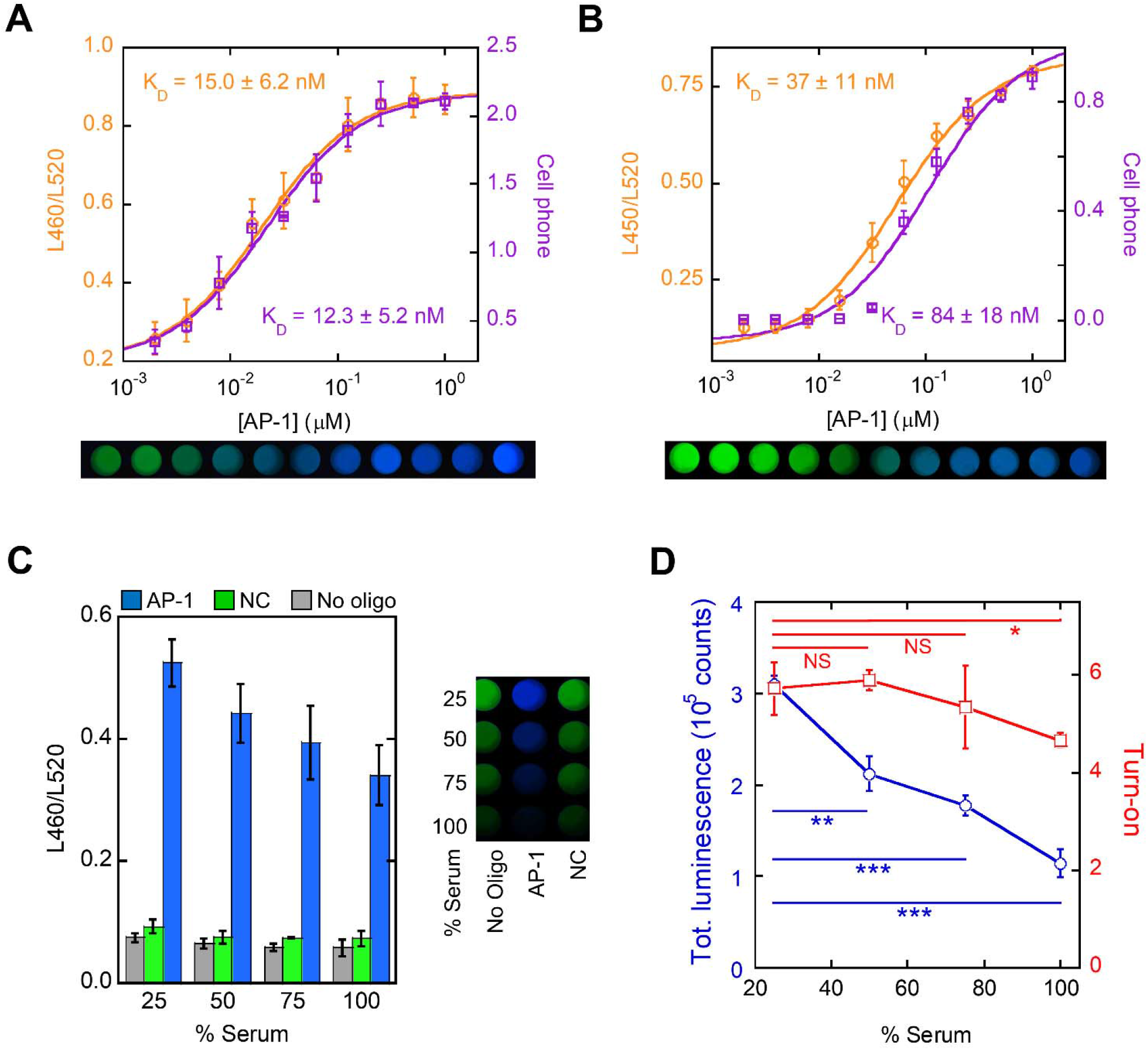
AP-1 binding activates nLuc-AFF in buffer and serum. Sensor turn-on by AP-1 binding in buffer containing **(A)** 0.1 mg/ml BSA or **(B)** 10 % FBS was quantified by ratio of luminescence at 460 nm and 520 nm (L460/520; orange) and ratio of blue to green channel intensities from cell phone images (purple). Solid lines represent best fits to the one-site quadratic binding equation. K_D_ values are average ± s.d. (n = 3 – 4). **(C)** nLuc-AFF turn-on as a function of serum concentration was quantified by L460/L520 (left) and cell phone images (right). AP-1 activated the biosensor, but NC oligo did not. Error bars are s.d. (n = 3). **(D)** Total biosensor luminescence decreased with serum content (blue), but fold turn-on (calculated by ratio of L460/L520 values in the presence and absence of AP-1) remained similar. Lines are meant to guide the eye only. Error bars are s.d. (n = 3) and p-values are from unpaired t-test. *p<0.05; **p<0.01; ***p<0.001; ****p<0.0001; NS, not significant.

It is important for a biosensor to function in dirty environments. This is especially problematic for luciferase-based sensors, as many existing designs show diminished luminescence in serum due to absorbance by extraneous components^24^. In 10 % serum, nLuc-AFF bound to AP-1 with similar affinity (K_D_ = 37 nM ± 11 nM; Fig. 2B) as in buffer, as determined by plate reader and cell phone camera. As serum concentration was raised to 100 %, we observed a decrease in overall luminescence intensity (Fig. 2D), as expected from serum absorbance, but the change in blue:green ratio due to biosensor activation remained constant within error (Fig. 2C, Fig. 2D). Raw cell phone images confirm that the samples are blue in the presence of AP-1 and green in the presence of nonconsensus (NC) DNA (Fig. 2C). This result, and the close agreement between the plate reader and cell phone camera data (Fig. 2A, Fig. 2B), demonstrate that it is possible to quantify the turn-on response using only a cell phone camera, by simply dividing the intensities of the blue and green channels without any image processing.

### Compatibility with toehold mediated strand displacement

Having developed the nLuc-AFF biosensor that is activated by AP-1, our next goal was to couple presentation of the AP-1 sequence to binding of other DNA sequence inputs. Toehold mediated strand displacement (TMSD)^25,26^ is a powerful tool that has been extensively used in applications such as signal amplification^6,7^, RNA computation^8,9^, and many others. It relies on the ability of an invader strand (ssDNA or ssRNA) to hybridize to a short ssDNA (the toehold) that overhangs from a dsDNA stem-loop structure in which the stem and toehold comprise the sequence complementary to that of the invading strand. The invading strand then fully hybridizes with the dsDNA hairpin, displacing the strand of identical sequence to the invader.

We selected a 24-nt sequence from the SARS-CoV-2 (SARS2) genome to serve as the invading strands for proof-of-concept. We designed two DNA hairpins in which the 5’ stem and the 5’ toehold consisted of the sequence complementary to the SARS2 oligonucleotide. The hairpins differed only by having the single-stranded AP-1 sequence (ssAP-1) (probe 1) or its complement (probe 2) embedded in their loops. Binding of the SARS2 oligonucleotide opens both probes, exposing their loops. The ssAP-1 sequences then hybridize, generating the activating complex that contains the duplex AP-1 input ligand for nLuc-AFF (Fig. 3A). Since AP-1 is a palindromic sequence with a single mismatch in the center, NUPACK^27^ simulations predicted that including ssAP-1 in the loop would extend the stem and make it too stable to be opened by the invading strand (Supporting Fig. S5A). We therefore added a 3-nt ‘bulge’ spacer between the stem and the ssAP-1 loop (Supporting Fig. S5B). The bulge was predicted to interrupt base stacking at the stem-loop junction, increasing efficiency of hairpin opening.

**Figure 3.**
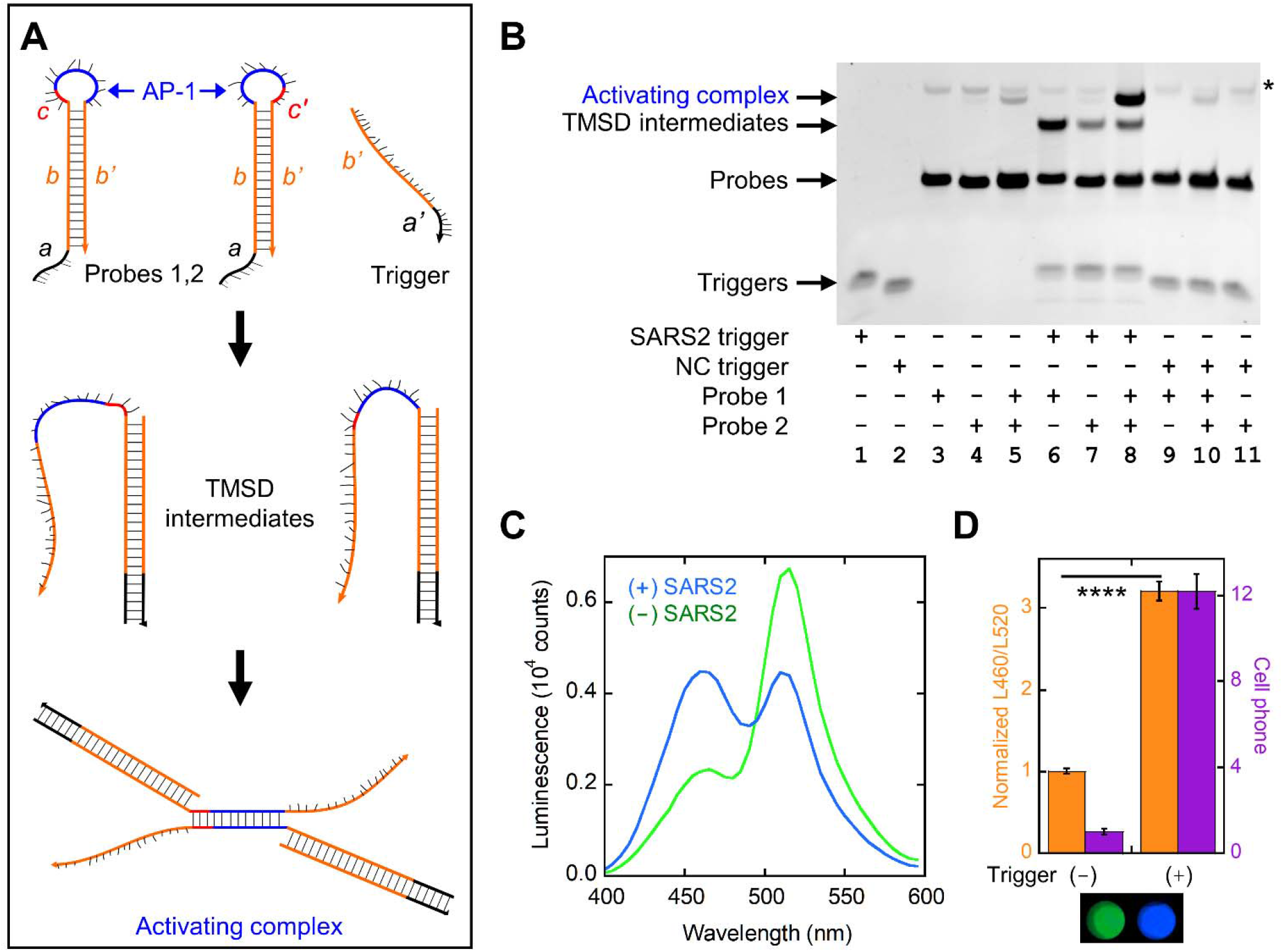
TMSD links an arbitrary DNA sequence to nLuc-AFF switching. **(A)** For detecting a desired ssDNA or RNA sequence (trigger), two DNA hairpins (probes 1 and 2) were designed with the trigger sequence comprising their stems (*b*) and toeholds (*a*) and the ssAP-1 sequences (blue) embedded in their loops. Probe 1 and probe 2 contain complementary ssAP-1 and the bulge *c*/*c’* sequences (see text) in their loops but are otherwise identical. The trigger opens both probes to form the TMSD intermediates, and the ssAP-1 sequences then hybridize to generate the activating complex. **(B)** Non-denaturing PAGE detected an intense band of activating complex when probe 1, probe 2, and CoV-2 trigger were mixed (lane 8). A faint band of activating complex was observed in mixtures of probe 1, probe 2, and NC trigger (lane 10) and probe 1 and probe 2 with no trigger (lane 5). The asterisk indicates an impurity that was most likely probe 1 dimer or probe 2 dimer. **(C)** nLuc-AFF mixed with probe 1 and probe 2 underwent a green-to-blue luminescence shift in the presence of CoV-2 trigger. **(D)** The CoV-2 trigger sequence turned on nLuc-AFF by 3.2-fold and 12.4-fold as quantified by L460/L520 ratio and cell phone images, respectively. The L460/L520 values were normalized to the spectra with no added trigger. Error bars are s.d. (n = 3). Unpaired t-test was performed using the L460/L520 biosensor response. ****p<0.0001.

We evaluated the performance of the designed hairpins by means of native PAGE. When the SARS2 oligonucleotide was incubated with either probe 1 or probe 2 alone, a new product of higher MW appeared, corresponding to the expected dimer of invading strand and opened probe (Fig. 3B). We observed a new, larger MW band upon adding the SARS2 oligonucleotide to both probes, consistent with formation of the activating complex.

Biosensor activation was tested by mixing nLuc-AFF with the two probes in the presence or absence of SARS2 oligonucleotide. We detected a decrease in green luminescence and an increase in blue luminescence only in the presence of the SARS2 strand (Fig. 3C). L460/L520 revealed a 3.2-fold change in biosensor turn-on along with a color change visible by cell phone camera (blue:green ratio change of 12.2) (Fig. 3D). This finding, along with the PAGE data, verify that the activating AP-1 duplex is formed from two hairpins only in the presence of the initiating SARS2 sequence. These results demonstrate that our biosensor can be used with TMSD-based DNA devices.

### Compatibility with DNA-based logic devices

To demonstrate the adaptability of our system, we applied it as a logic gate to process two ssDNA inputs. The first input strand (S1) was the same SARS2 oligonucleotide in Fig. 3 and the second input strand (S2) was identical except it bore a different 24-nt SARS2 genomic sequence. Using the approach described above, we then designed computational DNA probes to serve as logic gates, turning on via TMSD if: (i) either S1 or S2 are present (S1 OR S2 condition); (ii) both S1 and S2 are present (S1 AND S2 condition); (iii) S1 is present and S2 is not present (S1 NOT S2 condition). Input and probe oligonucleotides are shown in Fig. 4.

**Figure 4.**
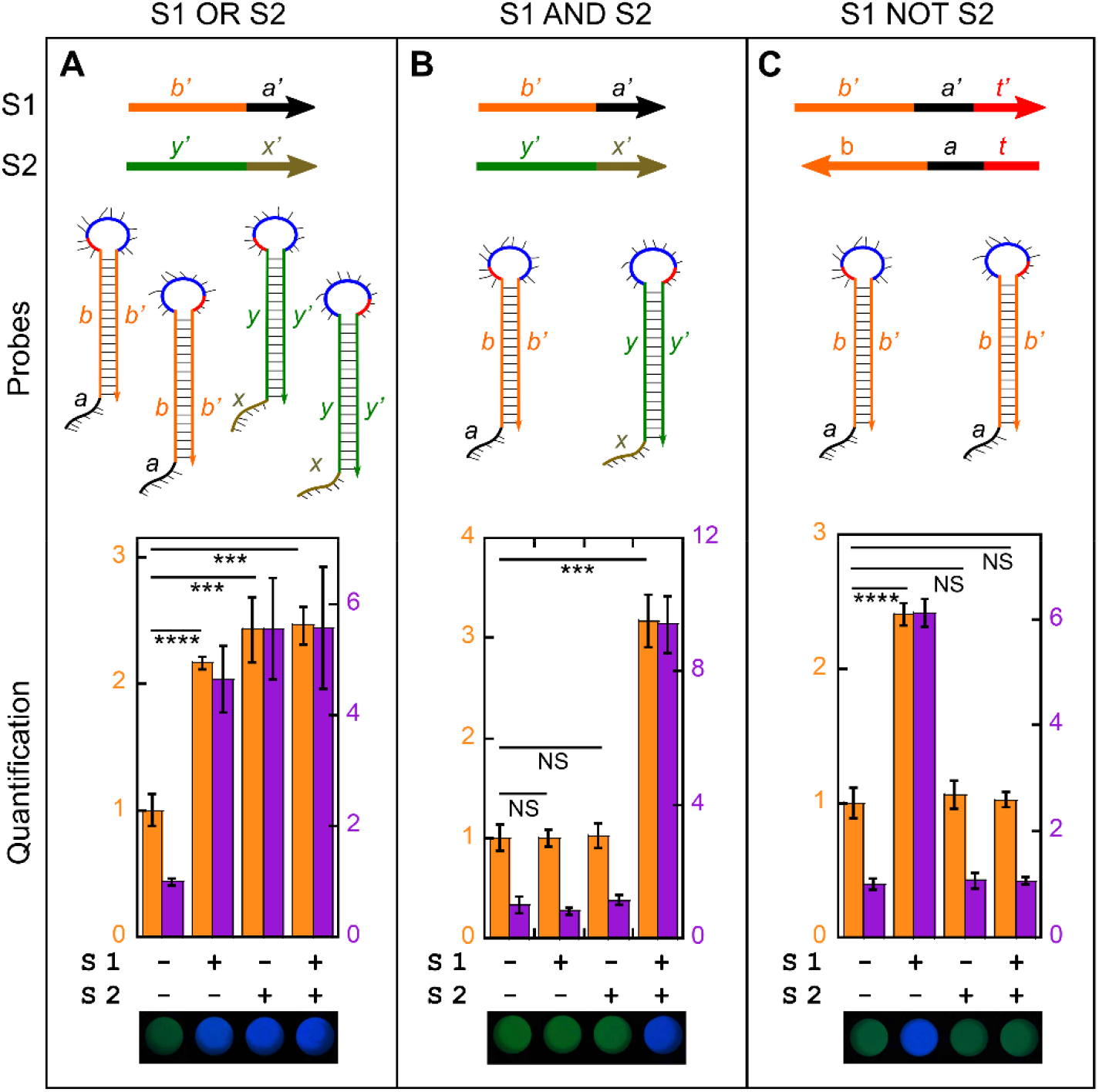
nLuc-AFF provides luminescent output for DNA computation. Input DNA strands, computational probes, and biosensor output are shown for (A) OR, (B) AND, and (C) NOT logic gates. The two input ssDNA strands, S1 and S2, were composed of one of two SARS2 genomic sequences (*b/b’* or *y/y’*) and a toehold (*a/a’* or *x/x’*). For the NOT condition, the S1 toehold was extended by 6-nt (*t’*) and S2 was an exact complement of S1. The additional 6-nt helps clamp S1 and S2 together, reducing fraying of the toehold in the S1/S2 complex that would result in probe invasion and nonspecific turn-on of the sensor^3^. The computational DNA probes were hairpins consisting of the SARS2 recognition sequences in their stems, complementary ssAP-1 sequences (blue) and bulge oligonucleotides (red) in their loops, and toehold sequences extending from the stems. These probes were kept constant in all four conditions of the logic gates. S1 and S2 were mixed in all four combinations and S1 OR S2, S1 AND S2, and S1 NOT S2 output was established qualitatively by cell phone camera (bottom images) in which green and blue indicate OFF and ON states, respectively. Sensor response was quantified by L460/L520 (orange bars) and by ratio of blue and green channels in the cell phone image (purple bars). Data were normalized to the condition where neither S1 nor S2 were present. Error bars are s.d. (n = 3). Unpaired t-tests were performed using the L460/L520 biosensor response. ***p<0.001; ***p<0.0001; NS, not significant.

The S1 OR S2 logic gate (Fig. 4A) was composed of two sets of hairpin pairs, one recognizing S1 and the other recognizing S2, each independently capable of opening and generating a separate activating complex. In agreement, we observed a large color shift when either of the two input strands were added to the biosensor-hairpin mix (Fig. 4A; Supporting Fig. S6A). The S1 AND S2 gate (Fig. 4B) contained a single probe that recognizes S1 and another that recognizes S2, such that both SARS2 sequences must be present to generate the activating complex. As expected, the biosensor was not activated by either S1 or S2 alone, and addition of both inputs gave rise to a large color shift (Fig. 4B, Supporting Fig. S6B). The S1 NOT S2 logic gate (Fig. 4C) consisted of a pair of hairpins that both recognize S1 and generate the activating complex upon binding. S2 was designed to base pair with S1 and thus inhibit S1 from initiating TMSD. S2 alone did not activate the biosensor-hairpin mix, and S1 alone gave rise to a large spectral shift, both as predicted (Fig. 4C, Supporting Fig. S6C). When we added S1 and S2, the blue:green ratio diminished to the same value that was observed without any inputs, satisfying the S1 NOT S2 condition.

### Compatibility with DNA aptamers

Sensing analytes of biological or clinical interest is a major goal of biosensor development. Many protein-based switches, including nLuc-AFF, employ natural, evolved, or *de novo* designed protein domains to bind the analytes and activate the output domains through allosteric mechanisms. This approach is powerful, but it is limited by the dual challenge of generating new binding domains that recognize the target of interest as well as undergo a large conformational change that transmit the binding signal to the output domain. DNA- and RNA-based aptamers provide a solution to the first challenge. Aptamers with high affinity and specificity for an arbitrary analyte can be obtained with relative ease by SELEX-based screening of large DNA libraries. The aptamers can then be derivatized with chemical dyes and molecular beacons for fluorescent output^28^ or attached to surfaces for detection by field-effect transistor devices^29^. It is not obvious, however, how the binding readout of aptamers can be expanded to include biological activities provided by protein and enzyme-based output domains.

Our nLuc-AFF system addresses the above limitation by acting as a ‘universal’ adapter that connects aptamer/analyte binding to bioluminescence. It is universal in the respect that it is designed to work with existing aptamers without any changes to nLuc-AFF or chemical derivatization/surface attachment of the aptamer. The only modification is addition of an oligonucleotide sequence to either the 5’ or 3’ end of the aptamer. To test that assertion, we sought to create a luminescent biosensor for serotonin using an aptamer that was developed elsewhere^29^. We modified the serotonin aptamer (green and red in Fig. 5A) by adding to its 5’ end a sequence consisting of ssAP-1 (blue) and an additional ‘clamp’ sequence (pink) that is complementary to first 8 nt of the aptamer (red). In the absence of ligand, the aptamer structure is unstable, and the red sequence base-pairs to the clamp, which, along with the blue sequence, forms the stem of a stable hairpin that prevents ssAP-1 from activating the biosensor (Fig. 5A, left structure). When the aptamer binds serotonin, it folds and reclaims the red sequence, opening the hairpin and exposing the ssAP-1 sequence in the loop (Fig. 5A, middle structure). This is essentially the DNA analog of the AFF mechanism. A second ssDNA oligonucleotide (naked AP-1) consisting of the sequence complementary to ssAP-1 and a truncated clamp (4-nt) is added to generate duplex AP-1 and activate nLuc-AFF (Fig. 5A, right structure). NUPACK was used to determine the optimal clamp lengths in the modified aptamer (to form a stable hairpin in the absence of ligand) and in naked AP-1 (to minimize false activation of the aptamer) (Supporting Fig. S7A).

**Figure 5.**
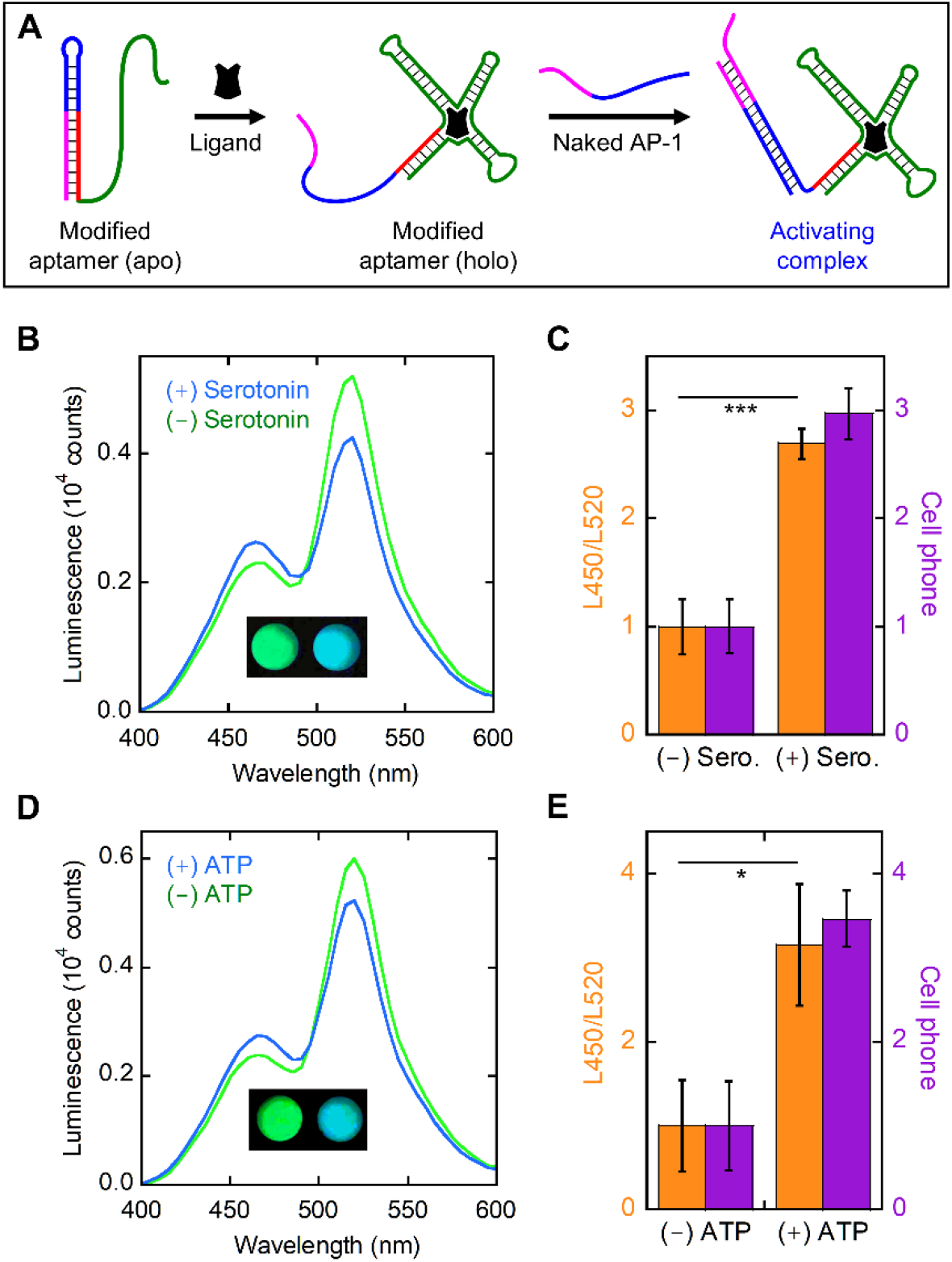
Small-molecule activation of nLuc-AFF via DNA aptamers. **(A)** The parental DNA aptamer (green and red) was modified by appending it to the ssAP-1 sequence (blue) and a ‘clamp’ sequence (pink) that’s complementary to the red sequence (left structure). Pairing of the pink and red strands stabilizes ssAP-1 in a hairpin structure, rendering it incapable of activating nLuc-AFF. Ligand binding induces the aptamer to fold, which displaces the clamp sequence and disrupts the ssAP-1 hairpin (center structure). The exposed ssAP-1/clamp sequence is then able to pair with an oligonucleotide consisting of the complementary ssAP-1 sequence and a portion of the clamp (‘naked AP-1’; blue and pink strand). The resulting complex (right structure) contains the duplex AP-1 sequence that activates the biosensor. **(B, D)** Mixing 10 μM serotonin and 2 mM ATP with their respective aptamers, naked ssAP-1, and nLuc-AFF triggered a green-to-blue luminescence change as detected by emission spectra and raw cell phone pictures (insets). **(C, E)** Quantifying sensor response by L450/L520 or ratio of blue/green channels from cell phone images revealed 2.7- and 3.0-fold turn-on (respectively) for the serotonin aptamer and 3.2- and 3.5-fold turn-on (respectively) for the ATP aptamer. For this calculation, signal from the biosensor with no aptamers or ligands present was subtracted from the observed signal, and the resulting values were normalized to the control with no ligand present. Unpaired t-tests were performed using the L460/L520 biosensor response. *p<0.05, ***p<0.001

We used native PAGE to establish that the modified aptamer did not bind naked AP-1 in the absence of serotonin. The aptamer alone ran as an intense, faster migrating band and a faint, slower migrating band (Supporting Fig. 7B). The slower species is likely a dimer, which is expected due to its largely palindromic AP-1 sequence along with the hairpin that protects it. We did not observe any additional products when naked AP-1 was added to the aptamer in the absence of serotonin. With serotonin, a new band was observed that likely corresponds to the activating complex depicted in Fig. 5A.

Having determined that a 1:1 ratio of aptamer:naked AP-1 yielded optimal results (Supporting Fig. S8), we evaluated sensor performance by adding 10 μM serotonin to samples containing 250 nM aptamer, 250 nM naked AP-1, and 30 nM nLuc-AFF. We observed a 2.7-fold ratiometric change in L450/L520 (Fig. 5B, Fig. 5C) and no change when the sensor was mixed with serotonin in the absence of aptamer (Supporting Fig. S8D). The color change was visible by cell phone camera (Fig. 5B inset), with an average of a 3-fold biosensor activation (Fig. 5C).

We next asked if our strategy was generalizable to other aptamers. We modified a well-characterized ATP aptamer^30^ to contain the ssAP-1 and clamp sequences using the approach described above, except we appended those nucleotides to the 3’-end of the ATP aptamer. As with the serotonin aptamer, we detected a spectral shift upon addition of 2 mM ATP, with a 3.1-fold increase in L450/L520 and a 3.6-fold change from the cell phone image (Fig. 5D, Fig. 5E). These results suggest that our design can be readily applied to existing aptamers to generate new luminescent sensors for a variety of small molecules and proteins.

Detection of serotonin is of considerable clinical interest. In conditions such as heparin induced thrombocytopenia (HIT) and catecholamine secreting carcinoid tumors, accurate quantification of platelet and serum serotonin can help aid diagnosis. HIT is a severe hypercoagulable condition with up to a 30 % mortality rate^31^. Although other clinical tests such as anti-platelet factor 4 (PF4) antibody titer are routinely performed, sensitive diagnosis relies on serotonin release assay^32^ (SRA). However, most healthcare settings source it to other facilities, causing large turn-around times. Our method provides an alternative to current methods used to perform SRA in low resource settings. The apparent affinity of the aptamer-biosensor (0.37 μM +/- 0.07 μM; Supporting Fig. S9) suggests that for HIT, levels of serotonin in healthy (0.11 – 0.36 μM) and diseased (1.5 - 46 μM) patients^33^ can be distinguished. Moreover, SRAs can also be used to diagnose vaccine induced thrombocytopenia^34^.

## Discussion

In this study, we bridge the fields of protein engineering and DNA engineering by developing a protein-based biosensor that is activated by different DNA inputs. The same nLuc-AFF protein can be paired with different DNA technologies for use in several applications. TMSD converts an arbitrary nucleotide sequence into the AP-1 input for the sensor, for the purpose of detecting a DNA/RNA sequence of choice or creating DNA-based computational devices with luminescent readout. Aptamers extend the potential targets of the nLuc-AFF biosensor to small molecules, metabolites, and proteins.

We note that nLuc-AFF retains some green emission even when fully saturated with AP-1. This appears due to the presence of truncated, always-green proteins that are generated by proteolytic cleavage within the 6* – 50* sequence, which is unpaired in the native fold and thus expected to be relatively unstructured. These fragments likely co-purify with the full-length sensor by means of their dimerizing GCN4 domains. Efforts to remove these truncated products (which appear as green fluorescent bands by SDS-PAGE) were unsuccessful (Supporting Fig. S11).

There are two main drawbacks to our current design. First is the slow turn-on rate (t_1/2_ = 8.25 h ± 0.37 h, Supplemental Fig. 10A). Based on our experience with other AFF-modified proteins, slow switching is due to a large kinetic barrier to unfolding of the native frame. Introducing mutations to destabilize the native fold can accelerate the overall switching rate. Care must be taken to preserve the thermodynamic balance between native and CP folds, and this can be accomplished by introducing the mutation in the region shared by the two folds (residues 51-171; Fig. 1B) or placing identical mutations in each of the duplicated segments. Destabilizing mutations can often be predicted by structural inspection or computational methods, but these do not always accelerate unfolding rates. A more rational approach is to simulate the AFF conformational change using weighted ensemble methods and identify amino acids that, when mutated, destabilize the native and CP folds but not the transition state ensemble^35^. However, even though the switch is slow to reach completion, robust signal change could be seen within 1 h (Supporting. fig. 10B). The cell phone images also showed a slight change in color in this time frame (Supporting. fig. 10C, D), suggesting that it may be possible to use nLuc-AFF for rapid diagnosis in low resource settings.

The second limitation of our system arises from the palindromic nature of the AP-1 sequence that activates the sensor. Our methods incorporate the activating sequence into various DNA structures, where it is cryptic until the triggering event (TMSD or ligand binding to aptamer) exposes it for hybridizing with its complement. In the case of TMSD, the DNA structures are hairpins, and introducing the palindromic ssAP-1 sequence into the loop can affect their metastability. Moreover, an unprotected ssAP-1 sequence (such as that in the complementary activating strand in our aptamer experiments) can homodimerize, giving rise to nonspecific activation. What is needed to resolve this problem is a DNA binding protein that recognizes a non-palindromic sequence, has a reasonably long N-to-C distance (≥20 Å), and can be engineered to be unstable in the absence of DNA. Zinc-finger DNA binding domains may be able to serve this purpose^36,37^. It may also be possible to use SELEX methods to identify a ssDNA aptamer that binds specifically to an altogether different input domain, i.e., a small protein that meets the distance and stability requirements mentioned above. This aptamer sequence would take the place of AP-1 and eliminate the need for the complementary activating strand.

The nLuc-AFF switch developed here has several notable attributes. It retains the DNA binding affinity and luminescent properties of the parent GCN4 and nLuc proteins, respectively, and it is not falsely activated by decoy DNA sequences. Its ratiometric output enables quantification by cell phone, by simply recording an RGB image and dividing the raw intensity of the blue channel by that of the green channel. The dual-color output establishes an internal normalization of intensity that makes it possible to quantify cell phone images. Finally, nLuc-AFF stands out from other luciferase-based platforms in that it performs well in serum without needing a secondary luminescent protein to calibrate for signal loss, opening the door to its potential use in clinical samples.

## Conclusions

Nature has provided enzymes—the Cas family of nucleases—that become activated by specific DNA or RNA sequences. It has not yet been feasible to introduce this mode of regulation into other proteins or enzymes. We have introduced a mechanism by which nanoluciferase was converted to a ratiometric biosensor that changes colors in response to a specific DNA sequence. The underlying AFF approach has been successfully applied to numerous other proteins^10,15,38,39^, suggesting that it can be used to create enzymes whose functions are switched on by specific DNA or RNA inputs.

## Methods

### Gene construction and protein purification

All expression constructs here were cloned into pET25 vector, which contains a C-terminal 6x-His tag. GeNL/pcDNA3 (gift from Takeharu Nagai, Addgene plasmid # 85200; http://n2t.net/addgene:85200; RRID:Addgene_85200) was used to PCR out the GeNL gene which was inserted into the expression vector at the NdeI/XhoI sites along with a C-terminal 6xHis tag. The GCN4 gene was inserted into the GeNL gene (at amino acid position 50 – 51 of the nLuc amino acid sequence), and KpnI and NheI restriction sites were added at the 5’- and the 3’-ends (respectively) of the resulting nLuc-AFF gene by extension PCR. Variants of this nLuc-AFF gene used in this study were constructed by synthesizing primers with various linkers as well as a NotI restriction site and assembling the new gene via overlapping PCR. Genes were fully sequenced after cloning. Protein sequences are shown in Supporting Fig. S12.

Proteins were expressed in *E. coli* BL21(DE3) cells with isopropyl β-D-thiogalactopyranoside induction for 16 – 18 h at 18°C. Cell pellets were resuspended in 20 mM Tris (pH 7.5), 0.5 M NaCl, 15 mM imidazole, 0.1% Tween-20 and lysed with a small amount of lysozyme followed by sonication (4×20 s pulses on ice). Viscosity was reduced by adding MgSO_4_ to final concentration of 5 mM along with DNase I, and the soluble fraction was loaded onto a nickel-nitrotriacetic acid column (Bio-Rad), and proteins were purified according to the manufacturer’s protocol. Eluted proteins were dialyzed into 20 mM Tris (pH 8.5), 150 mM NaCl and further purified using Superose 6 Increase 10/300GL (Cytiva) size-exclusion column. For purity determination, proteins were run on 0.02% SDS polyacrylamide gels and imaged on a Sapphire Biomolecular Imager (Azure Biosystems) to detect fluorescence from the mNG domain, which remains native in these conditions. nLuc-AFF purity was judged to be ∼70 % by this method, with the remaining 30 % consisting of C-terminally truncated products (Supporting Fig. S11).

### Luminescence measurements and image processing

Luminescence was measured using a SpectraMax i3 plate reader (Molecular Devices) with the following settings: 150 μL samples, 1 mm read height, 400 – 600 nm scans with 5 nm intervals). Furimazine (Aobious, catalog #36569) was added to a final concentration of 50 μM, and this solution was aged for at least 10 min at room temperature prior to recording luminescence spectra. Biosensor performance was quantified by the ratio of luminescence intensity at 460 nm and 520 nm (L460/L520). Quantification of samples by luminescence spectra and cell phone camera was performed on the same samples in the same plate (Corning Costar 96-well assay plates, white polystyrene, round bottom).

Cell phone images were taken using a OnePlus 7 Pro with the following settings: ISO 200, f/1.6 and auto WB. For quantification, images were split into the blue and green channels using ImageJ, and intensity in each channel was quantified by manually drawing a circle inside each well and calculating the average intensity. The average intensity in the blue channel was then divided by the average intensity in the green channel.

### Biosensor performance characterization

The performance of nLuc-AFF biosensor was characterized by the turn-on as well as the binding affinity of GCN4 to its consensus AP-1 sequence. We purified the nLuc-AFF three independent times (biological repeats) and performed three technical repeats of each biological repeat. To determine the switching efficiency, we mixed 30 nM of nLuc-AFF with either 2 μM of AP-1 containing double-stranded oligonucleotide (AGTGGA<u>GATGACTCATC</u>TCGTGC), 2 μM of the non-consensus double-stranded oligonucleotide (GTTCCAGGTTAAGAAGTGCTCTCAGGGTGGCGCGGC), or no oligonucleotide. Buffer was 20 mM Tris (pH 8.5), 150 mM NaCl, 0.1 mg/ml BSA. Samples were mixed and incubated overnight, in the dark, at room temperature, and scanned/photographed the next day. Equilibrium binding experiments were performed in triplicate by mixing 30 nM protein with various concentrations of AP-1 (2-fold serial dilutions starting from 2 μM) and incubating them overnight, in the dark, at room temperature. Buffer was 20 mM Tris (pH 8.5), 150 mM NaCl, and either 0.1 mg/ml BSA or 10 % FBS. Cell phone images and luminescence spectra were recorded for all samples, and the signals were fit to the one-site quadratic binding equation to obtain K_D_.

For experiments in serum, 2 M Tris (pH 8.5) was added to freshly thawed fetal bovine serum to a final concentration of 20 mM. This solution was then diluted to the desired serum concentration in 20 mM Tris (pH 8.5), 150 mM NaCl. The DNA and protein concentrations were identical to those described above. For integrated intensity measurements, biosensor was diluted in either 25 %, 50 %, 75 %, or 100 % serum to a final concentration of 30 nM, and total luminescence was calculated by integrating the spectra from 400 – 600 nm.

### Oligonucleotide Design and Purification

Oligonucleotides were designed using NUPACK to ensure metastability as well as high efficiency of TMSD. The two sequences of the nCoV genome used in this study were chosen from the CDC RT-PCR Diagnostic Panel: sequence N1 (cagattcaactggcagtaaccaga) and sequence N2 (tcagcgttcttcggaatgtcgcgc). The serotonin and ATP aptamers were also designed using NUPACK. Synthetic oligonucleotides were ordered from Eurofins Genomics without purification and were purified in-house using urea-PAGE (see Supporting Fig. S13 for methodology and oligonucleotide sequences).

### Toehold mediated strand displacement and logic gate experiments

Hairpins were snap-cooled separately in 20 mM Tris (pH 8.5), 150 mM NaCl at a concentration of 3uM, by heating to 95 °C for 3 min and rapidly cooling to room temperature using a thermal cycler. For validation by PAGE, 500 nM of probes were mixed with 500 nM of triggering oligonucleotide in 20 mM Tris (pH 8.5), 150 mM NaCl and incubated at room temperature for 3 h. Samples were then loaded on 12% acrylamide:bisacrylamide (19:1) gel in 1x Tris-borate-EDTA (TBE) buffer and run in TBE for 40 min at room temperature. Gels were then stained with ethidium bromide and imaged. For logic gate experiments, hairpins and nLucAFF were mixed to a final concentration of 300 nM and 30 nM, respectively, in 20mM Tris (pH 8.5), 150 mM NaCl, 0.1 mg/ml BSA. Triggering oligonucleotides were then added to a final concentration of 600 nM and the reaction was allowed to proceed overnight at room temperature before luminescence was recorded.

### Validation of nLuc-AFF with DNA aptamers

Serotonin and ATP aptamers were purified and snap-cooled before use as described in Supporting Figure S13. For PAGE assays, 250 nM of aptamer was mixed with 500 nM of naked AP-1 in 20 mM Tris (pH 8.5), 0.15 mM NaCl, whereupon either 10 μM 5serotonin, 2 mM ATP, or DMSO vehicle (final concentration 0.05%) was added, and the reaction was allowed to proceed for 2 h at room temperature. Samples were then loaded on a 12% polyacrylamide gel and run in TBE buffer. Gels were stained with ethidium bromide and imaged. For luminescence assays, 250 nM aptamer, and 250 nM naked AP-1, and 30 nM nLuc-AFF were mixed, and either 10 μM serotonin or 2mM ATP was added. Samples were kept overnight at room temperature before recording luminescence spectra the next day. Buffer was 20 mM Tris (pH 8.5), 150 mM NaCl, 0.1 mg/ml BSA, 5 mM MgCl_2_.

### Serotonin Aptamer Equilibrium Binding

Serotonin aptamer and the corresponding naked AP-1 oligonucleotide were individually snap-cooled as described above and mixed with 50 nM nLuc-AFF at a final concentration of 250 nM each in 20 mM Tris (pH 8.5), 150 mM NaCl, 0.1 mg/ml BSA, 5 mM MgCl_2_. Serotonin concentrations were prepared by 2-fold serial dilution starting 40 μM. Samples were maintained at room temperature overnight. Luminescence was quantified by both L460/L520 ratio and cell phone image and data were fit to the one-site binding equation to obtain K_D_.

### Biosensor turn-on kinetics

Freshly thawed biosensor (100 nM) was either mixed with 1 μM AP1 oligonucleotide or the same volume of buffer in 20mM Tris (pH 8.5), 150 mM NaCl, 0.1 mg/ml BSA. At the indicated timepoints, 150 μL of the mix was transferred to a fresh tube, furimazine added to final concentration of 50 μM, and spectra recorded as described above. The luminescence change was fit to a single exponential function to obtain the half time. For samples photographed by cell phone, 1 μM AP-1 oligonucleotide was added to a solution of 100 nM nLuc-AFF in 20 mM Tris (pH 8.5), 150 mM NaCl, 0.1mg/ml BSA in separate tubes separated in time by 15 min. 60 min after mixing the first sample, furimazine was added to a final concentration of 50 μM, and the reactions were aged for 10 min before imaging.

## Associated Content

### Supporting information available

The following file is available free of charge.

Supporting_Information_Sekhon_and_Loh.pdf, containing 13 supporting data figures.

## Acknowledgements

We thank Jeung-Hoi Ha for discussions. This work was supported by NIH grant R01GM115762 to S.N.L.

## Abbreviations

AFF: alternate frame folding
CP: circular permutant
GeNL: green enhanced nanolantern
mNG: mNeonGreen
NC: nonconsensus
nLuc: nanoluciferase
SARS2: SARS-CoV-2
TMSD: toehold-mediated strand displacement
SRA: serotonin release assay
HIT: heparin induced thrombocytopenia

