## Supplementary Information for "Engineering Protein Activity into Off-the-Shelf DNA Devices"

**Contents:**

**Figure S1.** Luminescence spectrum of nLuc-AFF with full-length duplicated segment (1* – 50*) and 12 amino acid linker in the CP frame shows mostly blue emission.

**Figure S2.** Optimization of nLuc-AFF constructs.

**Figure S3.** Optimizing the response of nLuc-AFF by varying CP linker length.

**Figure S4.** Equilibrium binding of nLucAFF to AP-1 in the presence of (A) 0.1 mg/mL BSA or (B) 10 % FBS.

**Figure S5.** Addition of bulge nucleotides to the stem-loop junction in probe 1 and probe 2 facilitates binding of the trigger oligonucleotide via TMSD.

**Figure S6.** Representative raw luminescence spectra for logic gate conditions tested in Fig. 4.

**Figure S7.** Modifying serotonin aptamer for activating nLuc-AFF.

**Figure S8.** Optimization of ssAP-1 and serotonin aptamer concentrations.

**Figure S9.** Equilibrium binding isotherm of serotonin with the aptamer and naked AP-1 (250 nM each) in the presence of 50 nM nLuc-AFF.

**Figure S10.** Turn-on kinetics of nLuc-AFF after addition of AP-1 oligonucleotide.

**Figure S11.** Purity of nLuc-AFF judged by 0.02 % SDS-PAGE with mNeonGreen fluorescence detection.

**Figure S12:** Amino acid sequences of nLuc-AFF constructs used in this study.

**Figure S13.** Sequences of DNA oligonucleotides and purification protocol.

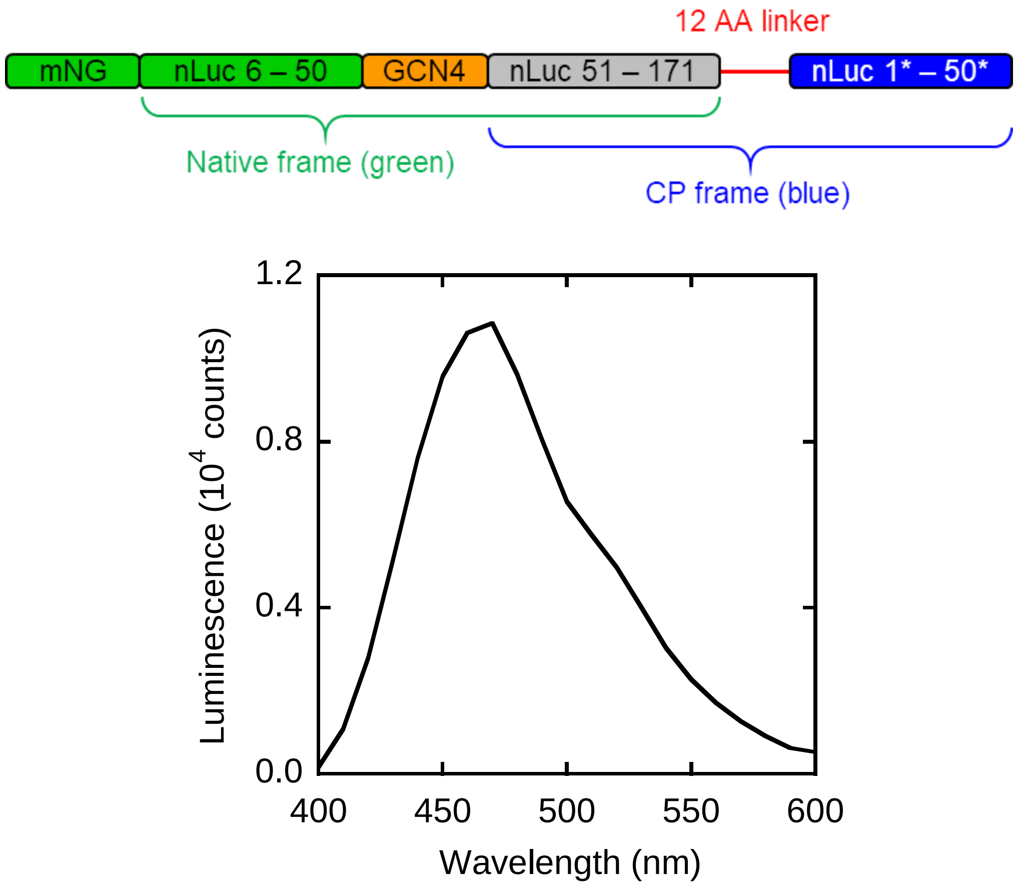


**Figure S1. Luminescence spectrum of nLuc-AFF with full-length duplicated segment (1* – 50*) and 12 amino acid linker in the CP frame shows mostly blue emission.**

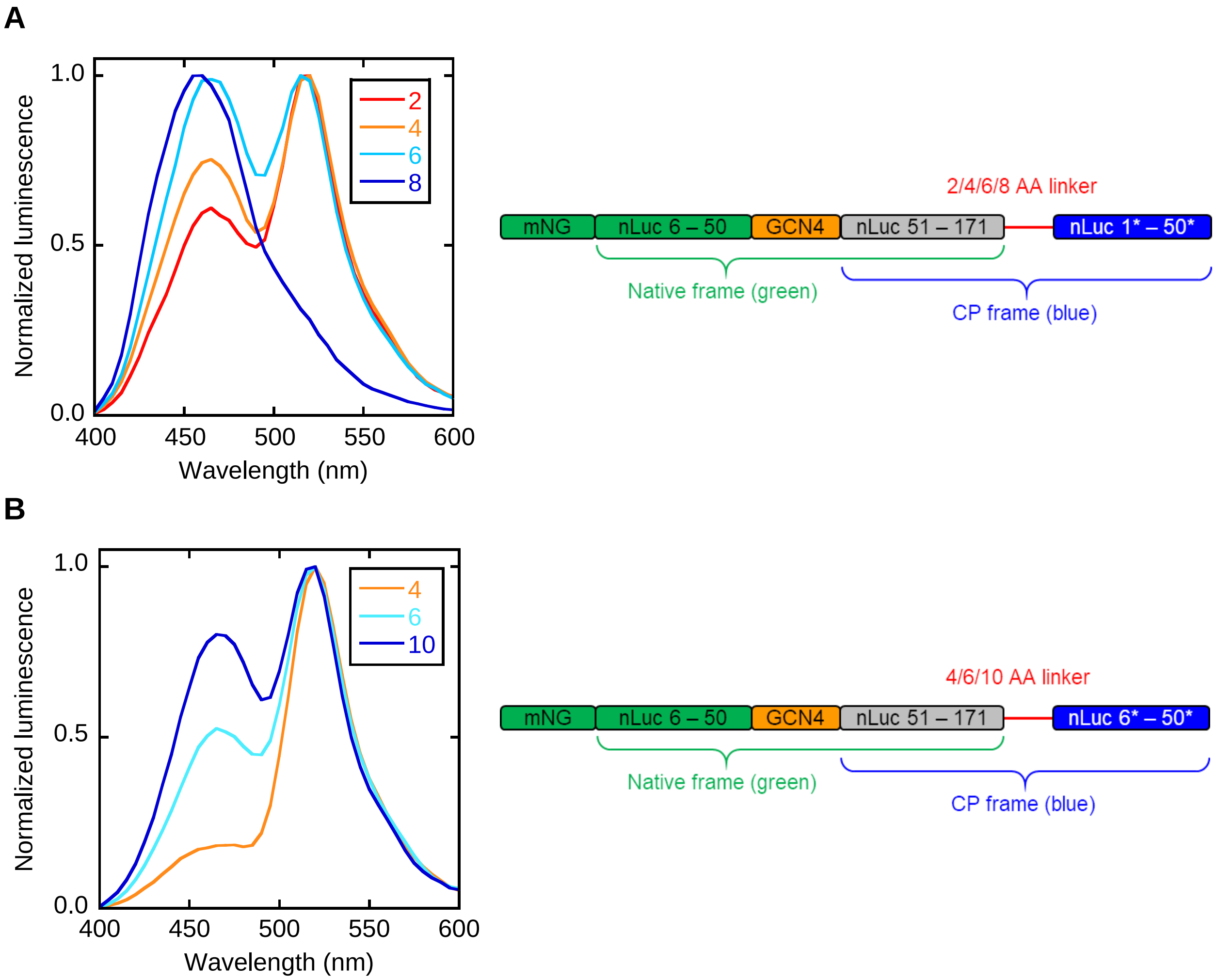


**Figure S2. Optimization of nLuc-AFF constructs. (A)** nLuc-AFF with full-length duplicated segment (1* – 50*) exhibits increasing populations of the CP fold with increasing CP linker length, as shown by the shift from green to blue luminescence. Insets next to luminescence spectra indicate the number of amino acids in the CP linkers. **(B)** nLuc-AFF with truncated duplicated segment (6* – 50*) show a trend similar to that in panel A, but with a stronger bias towards the native (green) frame. The scaffold shown in panel B was therefore chosen for further optimization.

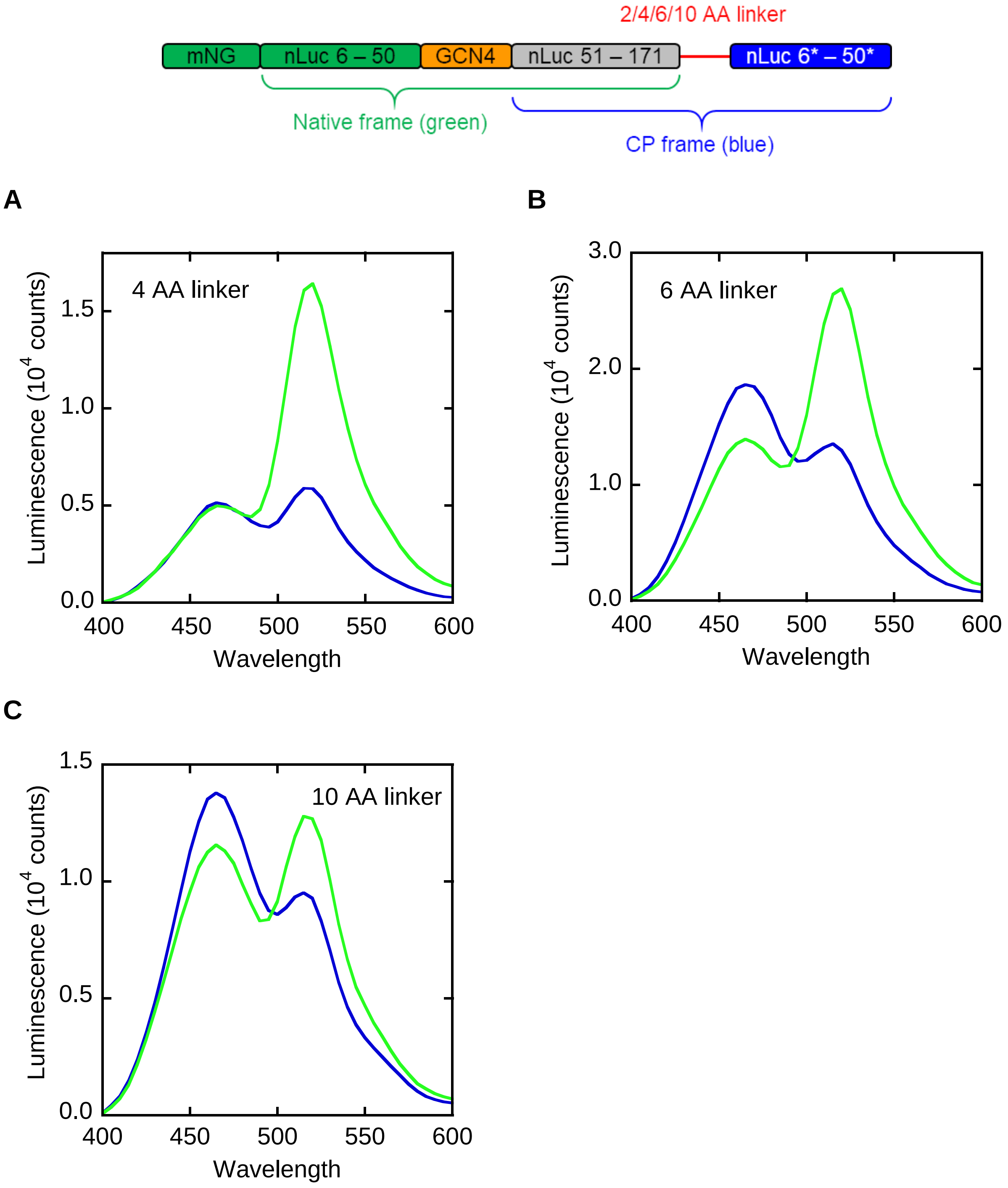


**Figure S3. Optimizing the response of nLuc-AFF by varying CP linker length.** The construct shown at top was engineered with **(A)** 4 AA linker, **(B)** 6 AA linker, and **(C)** 10 AA linker. The resulting sensors were mixed with AP-1 (blue spectra) or NC oligonucleotide (green spectra) and their luminescence spectra recorded. All sensor constructs exhibited a green-to-blue emission shift on addition of AP-1. The 6 AA linker in panel B showed the strongest response and was chosen for all further experiments.

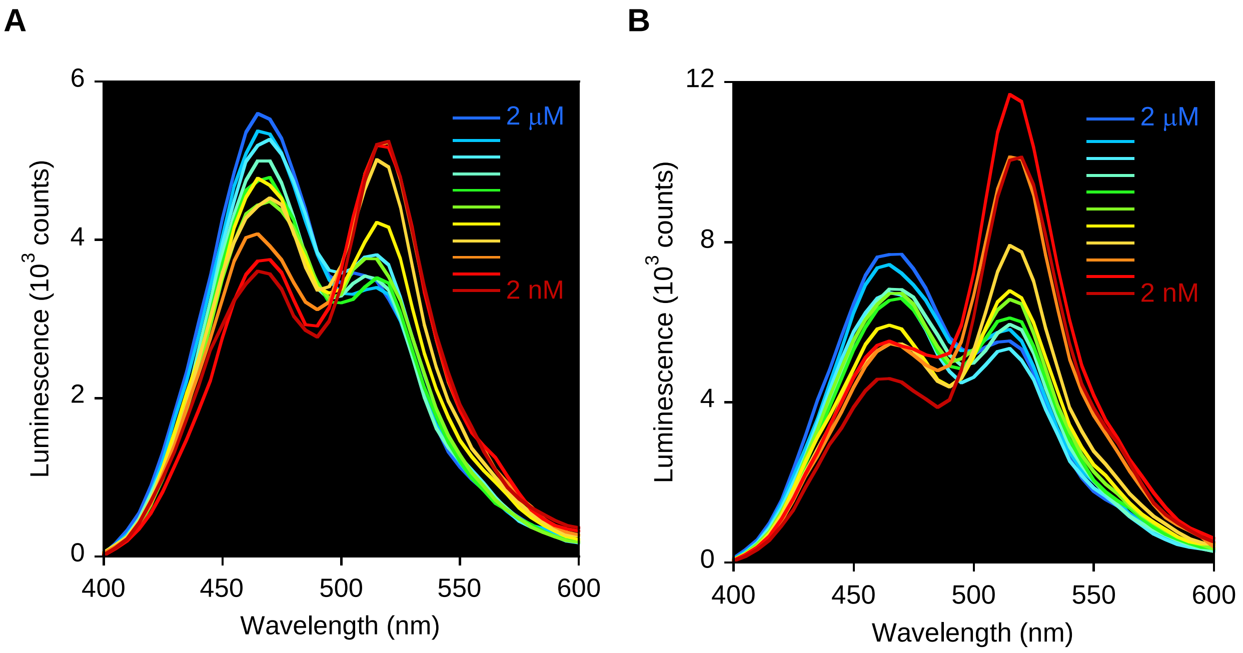


**Figure S4. Equilibrium binding of nLucAFF to AP-1 in the presence of (A) 0.1 mg/mL BSA or (B) 10 % FBS.** Insets indicate the concentration of AP-1 with each bar indicating a 2-fold change in concentration.

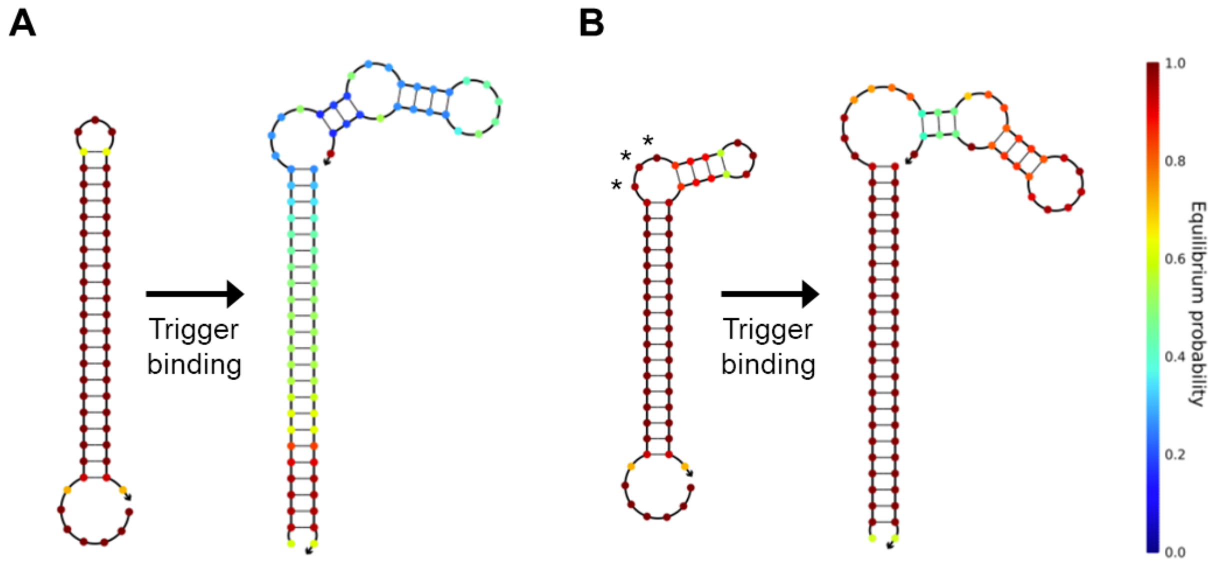


**Figure S5. Addition of bulge nucleotides to the stem-loop junction in probe 1 and probe 2 facilitates binding of the trigger oligonucleotide via TMSD.** The NUPACK-predicted structure of probe 1 (depicted schematically in Fig. 3A) is shown **(A)** without any bulge nucleotides, and **(B)** with three bulge nucleotides (asterisks). TMSD-mediated binding of the trigger produces the structures to the right of the arrows, with the trigger oligonucleotide comprising the right strand of the stem. Inclusion of the bulge nucleotides increases the efficiency of TMSD, as shown by the greater base pair probability of the probe-trigger complex in panel B compared to the probe-trigger complex in panel A.

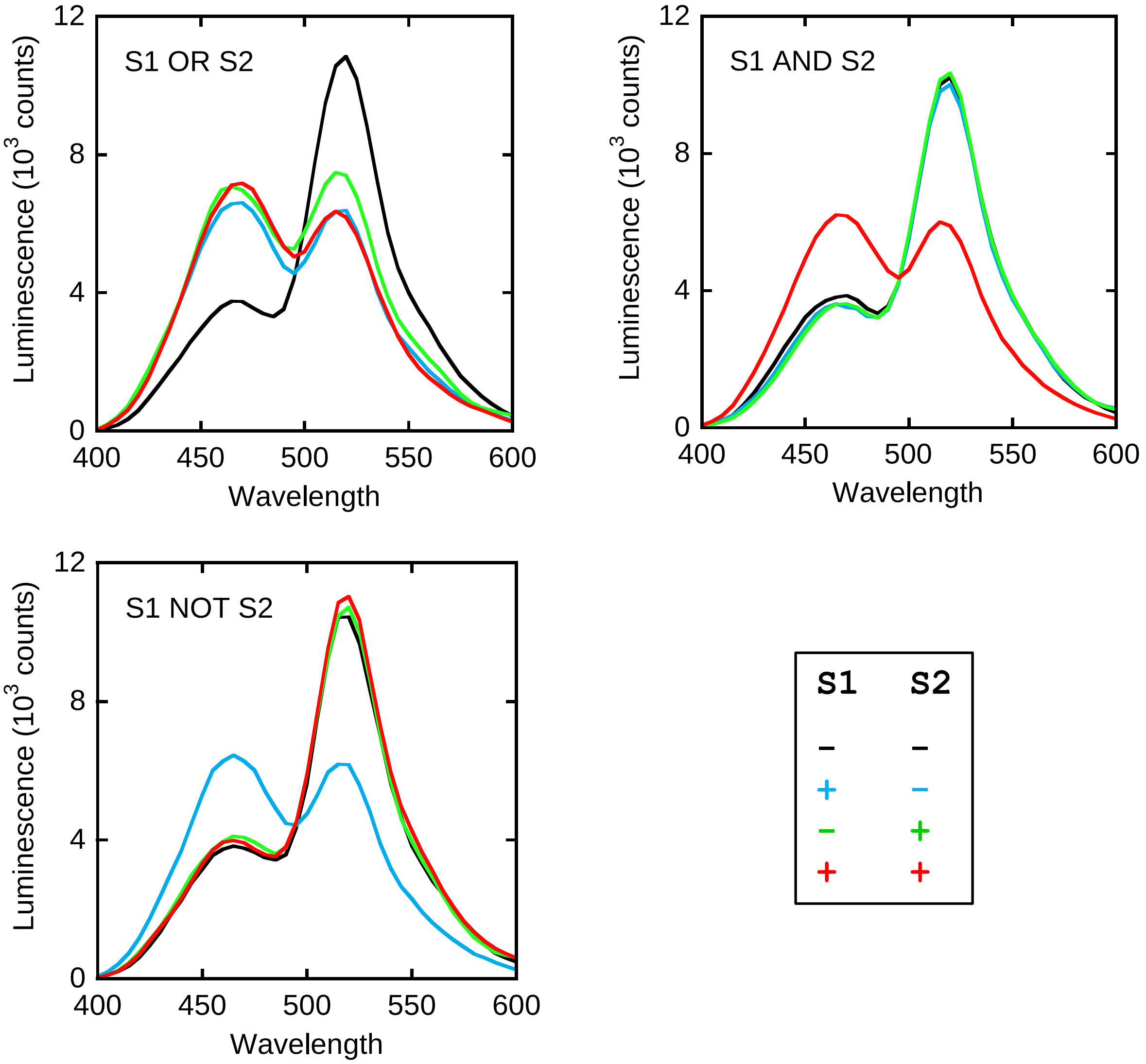


**Figure S6. Representative raw luminescence spectra for logic gate conditions tested in Fig. 4.** Colors of the spectra correspond to the mixtures of the S1 and S2 input strands indicated at lower right.

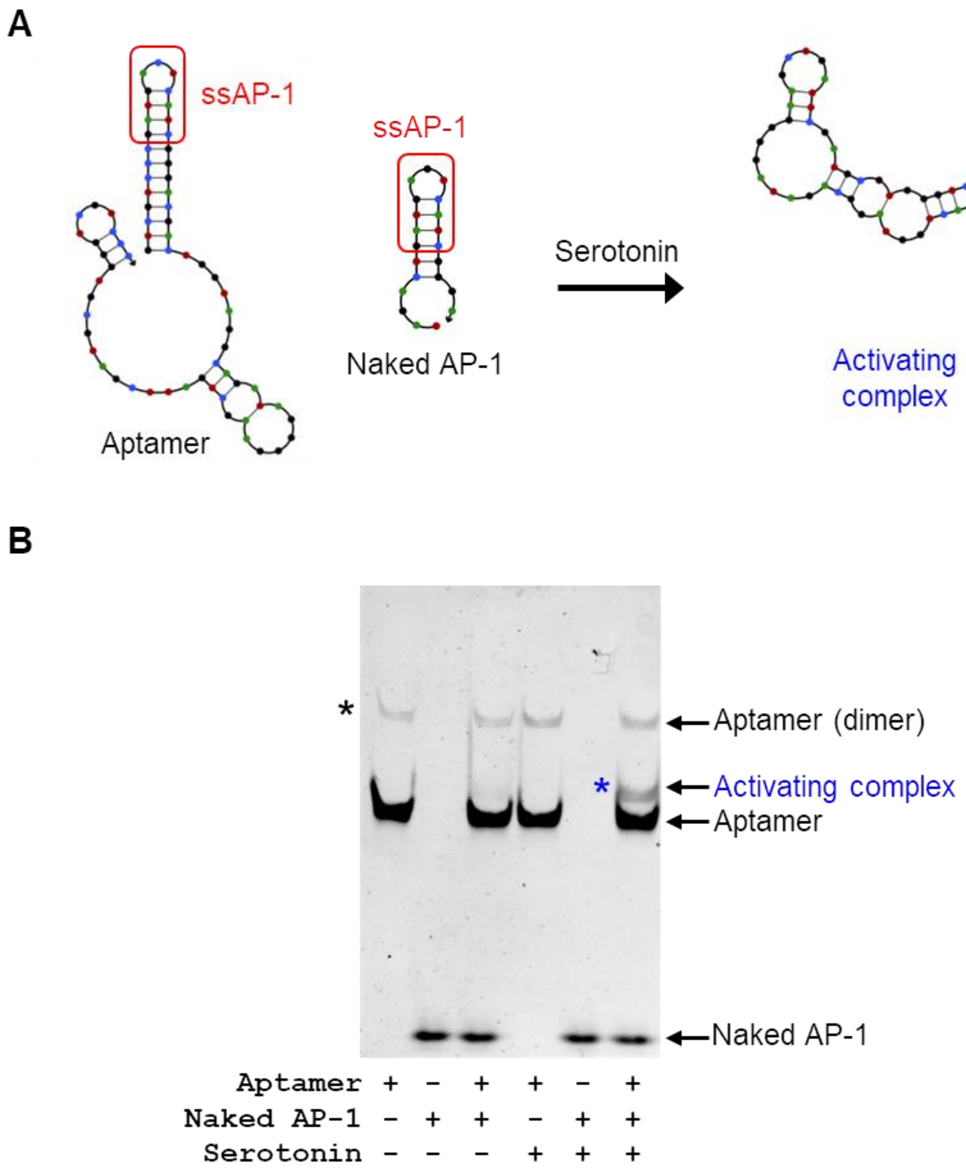


**Figure S7. Modifying serotonin aptamer for activating nLuc-AFF. (A)** NUPACK-predicted structures of the serotonin DNA aptamer with appended ssAP-1 and clamp sequences at the 5'-end (left structure), and the naked AP-1 oligonucleotide with complementary ssAP-1 and partial clamp sequences (middle structure). When the aptamer binds serotonin, it combines with the naked AP-1 oligonucleotide to generate the activating complex (right structure), with the full duplex AP-1 site shown in the red box. **(B)** Nondenaturing PAGE shows the appearance of a new band (the activating complex, blue asterisk) only in the presence of serotonin. Black asterisk indicates the species that likely represents a dimer of the aptamer.

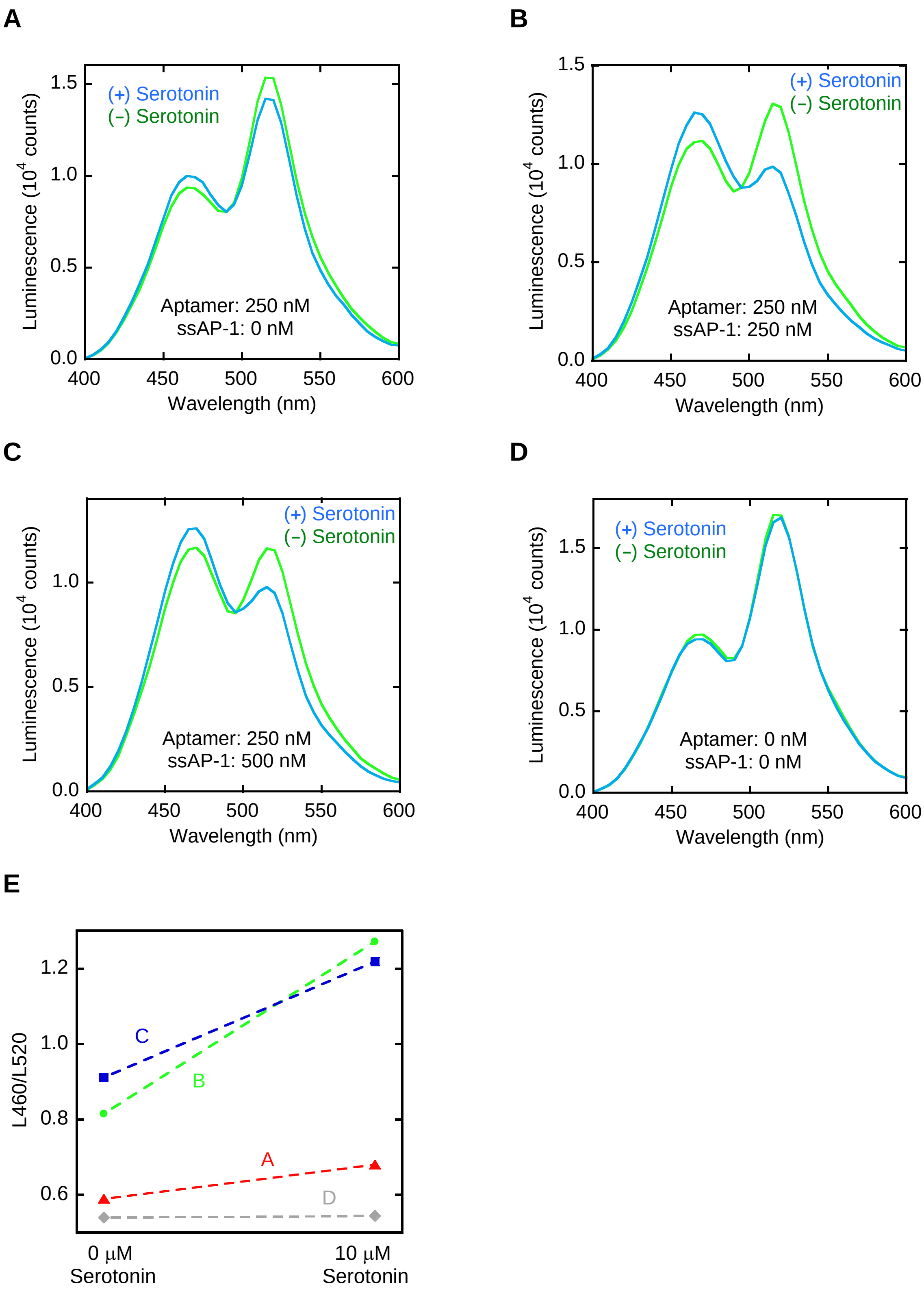


**Figure S8. Optimization of ssAP-1 and serotonin aptamer concentrations. (A – D)** Aptamer and ssDNA-1 were mixed at the indicated concentrations and added to nLuc-AFF.

Luminescence spectra were recorded with 10 μM serotonin (blue) or DMSO vehicle (green). **(E)** Luminescent color change (quantified by L460/L520) is shown for each of the data sets in panels A – D. Dashed lines are meant to guide the eye only. The ratiometric change in panel B (250 nM aptamer and 250 nM ssAP-1) was judged to be the best and these concentrations were used in subsequent experiments.

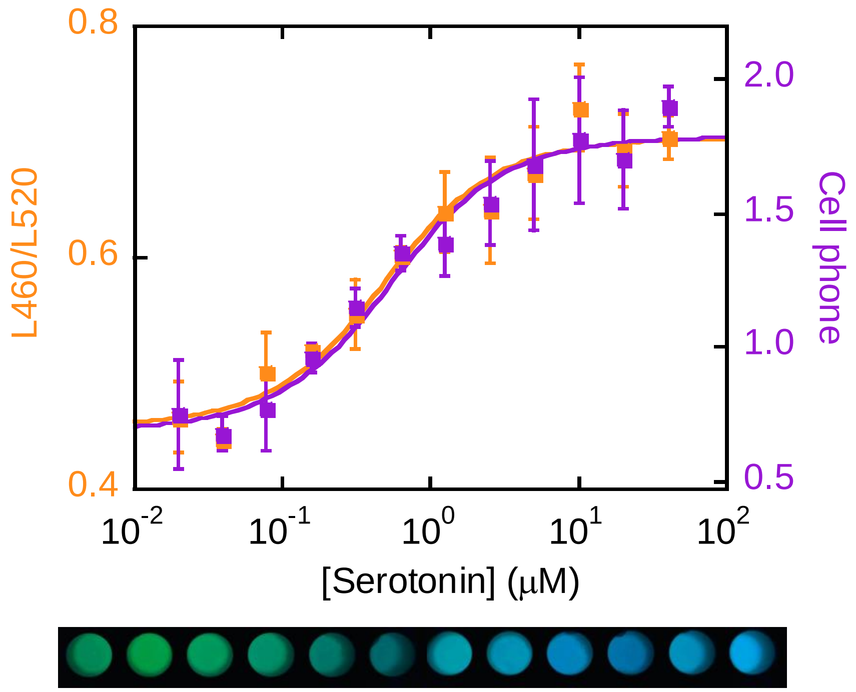


**Figure S9. Equilibrium binding isotherm of serotonin with the aptamer and naked AP-1 (250 nM each) in the presence of 50 nM nLuc-AFF.** Binding was monitored by L460/L520 from luminescence spectra (orange) and by the ratio of intensities in blue and green channels of cell phone images. Lines indicate best fits to the 1-site quadratic binding equation, with $K_D$ = 370 nM ± 70 nM (L460/520) and 418 nM ± 170 nM (cell phone camera). Error bars are s.d. (n = 3).

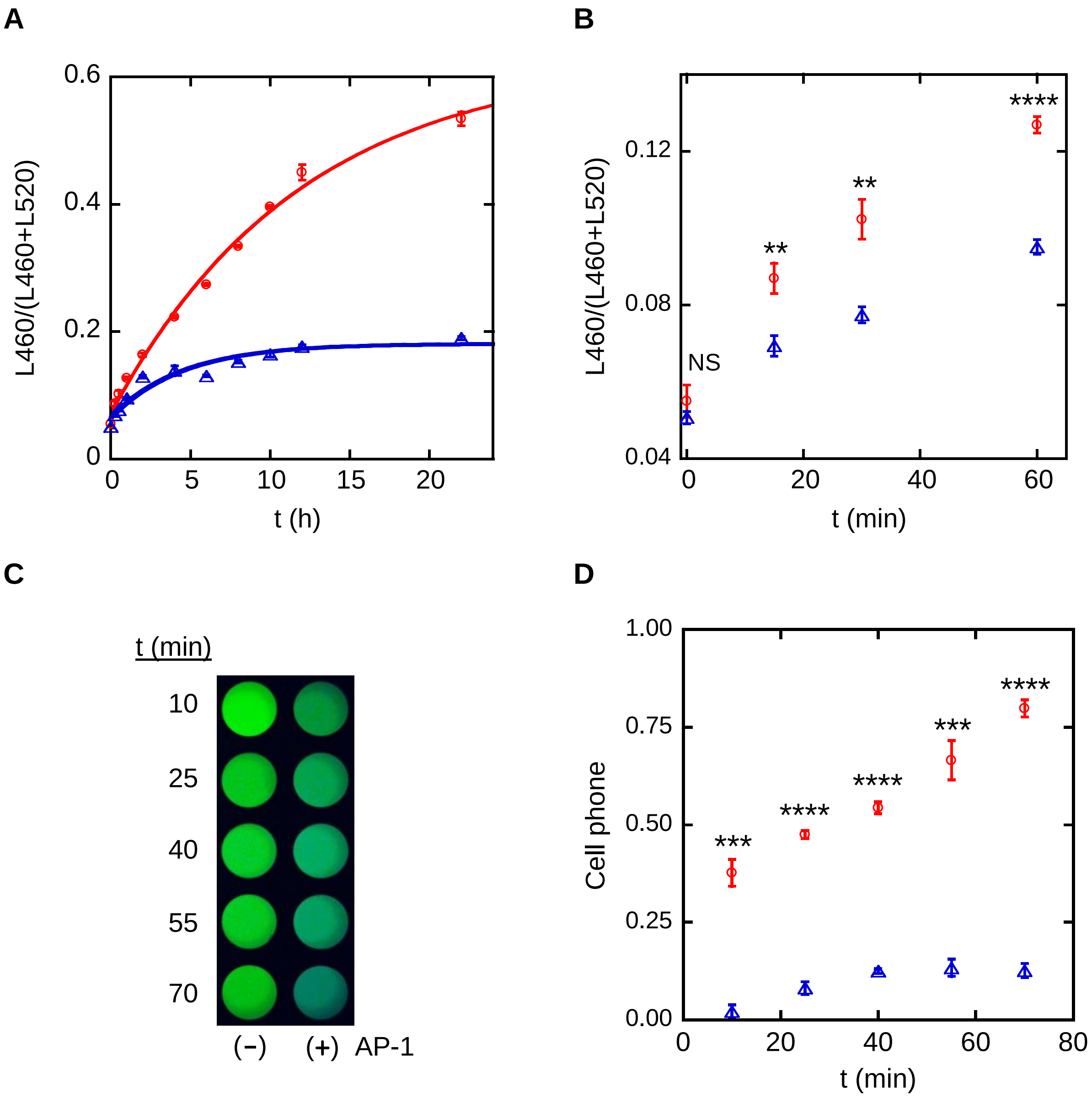


**Figure S10. Turn-on kinetics of nLuc-AFF after addition of AP-1 oligonucleotide. (A)** AP-1 activates nLuc-AFF with $t_{1/2}$ = 8.25 h ± 0.37 h (red line and circles). nLuc-AFF exhibits a small turn-on in the absence of AP-1 (blue line and triangles) that is caused by the shift in temperature from storage conditions (4 °C) to experimental temperature (~22 °C) with $t_{1/2}$ = 3.10 h ± 0.65 h. Lines are best fits to a 1-exponential function and error bars are s.d. (n = 3). **(B)** The same data in panel A are zoomed in to show that biosensor activation is significant (relative to the negative control) within one hour. **(C)** Raw cell phone images and **(D)** quantification of cell phone images by ratio of blue:green channel intensity reveal significant color changes in less than an hour of

AP-1 exposure. Error bars in panel D are s.d. ($n = 3$). **$p<0.01$, ***$p<0.001$, ***$p<0.0001$, NS = not significant.

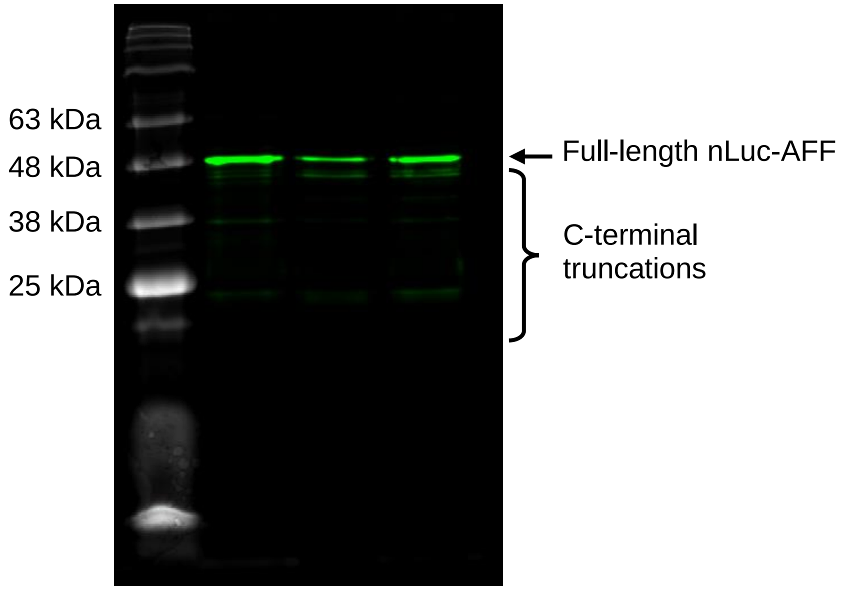


**Figure S11. Purity of nLuc-AFF judged by 0.02 % SDS-PAGE with mNeonGreen fluorescence detection.** The nLuc-AFF construct contains mNG and a 6x-HisTag fused to its N- and C-terminus, respectively. After purification by nickel-NTA chromatography, ~30 % of recovered fluorescent protein consisted of products that were smaller yet fluorescent, revealing that they were truncated at their C-termini. These impurities should not bind to nickel-NTA beads, but they likely dimerize to full-length nLuc-AFF via the GCN4 domains and co-purify. The truncated proteins likely account for the green background that remains at saturating concentrations of AP-1.

**(A)** nLuc-AFF:

MVSKGEEDNMASLPATHELHIFGSINGVDFDMVGQGTGNPNDGYEELNLKSTKGDLQFSPWILVPHIGYG
FHQYLPYPDGMSPFQAAMVDGSGYQVHRTMQFEDGASLTVNYRYTYEGSHIKGEAQVKGTGFPADGPVMT
NSLTAADWCRSKKTYPNDKTIISTFKWSYTTGNGKRYRSTARTTYTFAKPMAANYLKNQPMYVFRKTELK
HSKTELNFKEWQKAFTGFEDFVGDWRQTAGYNLDQVLEQGGVSSLFQNLGVSVTPIQRIVLSGEGTGGGS
SDPAALKRARNTEAARRSRARKLQRMKQLEDKVEELLSKNYHLENEVARLKKLVGERGASSGENGLKIDI
HVIIPYEGLSGDQMGQIEKIFKVVYPVDDHHFKVILHYGTLVIDGVTPNMIDYFGRPYEGIAVFDGKKIT
VTGTLWNGNKIIDERLINPDGSLLFRVTINGVTGWRLSERILAXXXEDFVGDWRQTAGYNLDQVLEQGGV
SSLFQNLGVSVTPIQRIVLSGENGLEHHHHHH

Color coding: mNeonGreen, nanoluciferase, GCN4 DNA binding domain, Linkers, Variable length CP linker (designated as XXX), HisTag.

**(B)** For Supporting Figure 2C, the XXX sequence consisted of AAVFTL, AASSVFTL, AAGASSVFTL, AASSGASSVFTL for the 2-, 4-, 6-, 8-AA CP linkers, respectively. VFTL is part of the WT nanoluciferase sequence.

**(C)** For Supporting Figure 2D, the XXX sequence consisted of AASS, AAGASS, AASSGASSVFTL for the 4-, 6-, 10-AA CP linkers, respectively.

**Figure S12: Amino acid sequences of nLuc-AFF constructs used in this study. (A)** Parent nLuc-AFF; **(B)** CP linker length variants with the nanoluciferase sequence VFTL included in the CP frame and counted as part of the CP linker; **(C)** CP linker length variants with the nanoluciferase sequence VFTL deleted from the CP frame.

**(A)** DNA sequences of oligonucleotides used in Figure 1 and Figure 2.

```
AP1          AGTGGAGATGACTCATCTCGTGC
AP1_rc       GCACGAGATGAGTCATCTCCACT
NC           GTTCCAGGTTAAGAAGTGCTCTCAGGGTGGCGCGGC
NC_rc        GCCGCGCCACCCTGAGAGCACTTCTTAACCTGGAAC
```

**(B)** DNA sequences of oligonucleotides used in Figure 3.

```
Trigger      cagattcaactggcagtaaccaga
Probe 1      tctggttactgccagttgaatctgatcGATGACTCATCcagattcaactggcagta
Probe 2      tctggttactgccagttgaatctgGATGAGTCATCgatcagattcaactggcagta
```

**(C)** DNA sequences of oligonucleotides used in Figure 4.

S1 OR S2 logic gate:

```
S1           cagattcaactggcagtaaccaga
S2           tcagcgttcttcggaatgtcgcgc
S1_Probe 1   tctggttactgccagttgaatctgatcGATGACTCATCcagattcaactggcagta
S1_Probe 2   tctggttactgccagttgaatctgGATGAGTCATCgatcagattcaactggcagta
S2_Probe 1   gcgcgacattccgaagaacgctgaatcGATGACTCATCtcagcgttcttcggaatg
S2_Probe 2   gcgcgacattccgaagaacgctgaGATGAGTCATCgattcagcgttcttcggaatg
```

S1 AND S2 logic gate:

```
S1           cagattcaactggcagtaaccaga
S2           tcagcgttcttcggaatgtcgcgc
S1_Probe 1   tctggttactgccagttgaatctgatcGATGACTCATCcagattcaactggcagta
S2_Probe 2   gcgcgacattccgaagaacgctgaGATGAGTCATCgattcagcgttcttcggaatg
```

S1 NOT S2 logic gate:

```
S1           cagattcaactggcagtaaccagattaacc
S2           ggttaatcagcgttcttcggaatgtcgcgc
S1_Probe 1   tctggttactgccagttgaatctgatcGATGACTCATCcagattcaactggcagta
S1_Probe 2   tctggttactgccagttgaatctgGATGAGTCATCgatcagattcaactggcagta
S2_Probe 1   gcgcgacattccgaagaacgctgaatcGATGACTCATCtcagcgttcttcggaatg
S2_Probe 2   gcgcgacattccgaagaacgctgaGATGAGTCATCgattcagcgttcttcggaatg
```

**(D)** DNA sequences of aptamers and naked AP-1 oligonucleotides used in Figure 5.

```
Serotonin aptamer:
gtcgtcccGATGACTCATCgggacgactggtaggcagataggggaagctgattcgatgcgtgggtcgtccc
Serotonin naked AP-1:
tagactGATGAGTCATCggga
ATP aptamer:
acctggggggagtattgcggaggaaggtGATGACTCATCaccttcct
ATP naked AP-1:
aaggtGATGAGTCATCa
```

**Figure S13. Sequences of DNA oligonucleotides and purification protocol.** Nucleotide colors are matched to those in the figures. The AP-1 sequence is capitalized and in blue. Synthetic oligonucleotides were purified using urea-PAGE (7 M urea, 8 % acrylamide:bis 19:1, 1x Tris-borate-EDTA). Samples (100 µL of 50 µM oligonucleotide in 50 % formamide, 0.01% bromophenol blue) were loaded on a 55 °C prewarmed 16.5 cm x 19 cm gel and run at 200 V for 2 h with heating by a circulating 55 °C water bath. Gels were stained in 2 % methylene blue for 20 m and briefly rinsed with water. Bands were excised, crushed, and mixed with ~5 volumes elution buffer (20 mM Tris pH 7.5, 0.5 M NaCl, 1 mM EDTA). After 2 freeze-thaw cycles at -80 °C, the tubes were shaken overnight at 37 °C. The samples were then centrifuged to remove gel fragments, and an equivalent volume of 1-butanol was added with vigorous vortexing. The organic phase was discarded, and 1/5 volume of 3 M sodium acetate (pH 5.2) was introduced along with 2.5x volume of 95 % ethanol pre-chilled at -80 °C. These samples were kept at -80 °C for 1 h then centrifuged at 16,000 x *g* for 30 m at 4 °C. The supernatant was discarded and the pellet was rinsed with chilled 95 % ethanol, centrifuged at 16,000 x *g* for 30 m at 4 °C, and dried at 37 °C for 20 min. The pellet was suspended in $ddH_2O$ and concentration was determined by nanodrop.
